# Exercise mitigates sleep-loss-induced changes in glucose tolerance, mitochondrial function, sarcoplasmic protein synthesis, and circadian rhythms

**DOI:** 10.1101/2020.06.21.163733

**Authors:** Nicholas J Saner, Matthew J-C Lee, Jujiao Kuang, Nathan W Pitchford, Gregory D Roach, Andrew Garnham, Amanda J Genders, Tanner Stokes, Elizabeth A Schroder, Karyn A Esser, Stuart M Phillips, David J Bishop, Jonathan D Bartlett

## Abstract

Sleep loss has emerged as a risk factor for the development of impaired glucose tolerance. The mechanisms underpinning this observation are unknown; however, both mitochondrial dysfunction and circadian misalignment have been proposed. Given that exercise improves glucose tolerance, mitochondrial function, and alters circadian rhythms, we investigated whether exercise may counteract the effects induced by inadequate sleep. We report that sleeping 4 hours per night, for five nights, reduced glucose tolerance, with novel observations of associated reductions in mitochondrial function, sarcoplasmic protein synthesis, and measures of circadian rhythmicity; however, incorporating three sessions of high-intensity interval exercise (HIIE) during this period mitigates these effects. These data demonstrate, for the first time, a sleep loss-induced concomitant reduction in a range of physiological processes linked to metabolic function. These same effects are not observed when exercise is performed during a period of inadequate sleep, supporting the use of HIIE as an intervention to mitigate the detrimental physiological effects of sleep loss.

## Introduction

The detrimental effects of sleep loss on glucose tolerance are now well-established, and insufficient sleep is a risk factor for the development of type 2 diabetes (T2D) (1). In fact, sleep loss is comparable with other more traditional risk factors that are associated with the development of T2D, such as physical inactivity (1). Several studies have shown that periods of sleep restriction, or reduced time in bed (TIB), typically with a sleep opportunity of 4 to 5 h per night, cause significant reductions in a range of indices related to glucose metabolism (2-5). The severity of this effect can be seen with only one night of either sleep restriction (4 h TIB) or sleep deprivation (e.g., no sleep), which can reduce insulin sensitivity (6-9). Despite these findings, there are limited data explaining the physiological and molecular changes that underpin these effects. As a large proportion of the population do not meet the current sleep recommendations (e.g., 7 to 9 h, per night) (10, 11) and inadequate sleep is a consequence of many occupations (12, 13), gaining a better understanding of these mechanisms may help to tailor specific interventions aimed at counteracting the detrimental effects of sleep loss.

The physiological mechanisms that underpin the impairment of glucose tolerance following sleep restriction are likely multifactorial. While not previously investigated in the context of sleep loss, the development of insulin resistance has been associated with a reduction in mitochondrial content and impaired mitochondrial respiratory function (14, 15). Furthermore, reductions in citrate synthase activity (a surrogate marker for mitochondrial content (16)) and mitochondrial respiratory function have also been reported in T2D patients, compared to obese non-diabetics, suggesting a link between mitochondrial changes and the development of insulin resistance and T2D (14, 15). Therefore, sleep-loss-induced reductions in glucose tolerance may, in part, be a consequence of changes in mitochondrial content, function, or the processes that regulate these properties - including mitochondrial dynamics and mitochondrial protein synthesis (17). In support of this, 120 h of sleep deprivation was associated with a 24% reduction in citrate synthase activity in human skeletal muscle (18). However, how these results translate to the context of the sleep loss commonly experienced in society, such as repeated nights of partial sleep loss, has not been determined and remains a critical gap in the literature.

The detrimental effect of sleep loss on glucose metabolism may also be associated with the misalignment of circadian rhythms (7, 8). One night of sleep deprivation (commonly experienced by 20% of the world’s population who perform shift-work) leads to a reduction in glucose tolerance and concomitant alterations in the expression of skeletal muscle clock genes (i.e., *Bmal1* and *Cry1* gene expression (7)) and the content of clock proteins (i.e., BMAL1) (8), which are known to regulate circadian rhythms at a molecular level (19). The functional significance of disrupting the molecular clock has been shown in genetic mouse models (i.e., Clock mutant mice and the *Bmal1* KO mouse), which display reduced glucose tolerance, mitochondrial respiratory function, and skeletal muscle contractile function (20, 21). However, the effect of sleep restriction on markers of circadian rhythmicity (e.g., skeletal muscle clock gene expression) and the potential implications of such changes have not previously been examined.

One approach to mitigating or ablating the impact of reduced sleep duration on glucose tolerance is via exercise (17). Regular endurance exercise has been shown to exert beneficial effects on glycaemic control via the activation of the insulin-independent signalling pathway (22). High-intensity interval exercise (HIIE) is a time-efficient format of endurance exercise, and is also a potent stimulus for the induction of mitochondrial biogenesis (23), with increases in sarcoplasmic and mitochondrial protein synthesis, and mitochondrial content and respiratory function, occurring concomitantly with improvements in glucose tolerance (24-27). This raises the intriguing hypothesis that exercise may also be useful to combat sleep-loss-induced impairments to glucose tolerance, which are not necessarily reversed by a period of recovery sleep alone (28-30). Furthermore, the same detrimental metabolic changes that occur in response to circadian misalignment and altered expression of clock genes may also be ameliorated by performing exercise (20, 31, 32). Consequently, HIIE may be able to mitigate the detrimental effects of sleep loss on glucose metabolism, by increasing mitochondrial content and function, and preventing changes in circadian rhythmicity (17).

Accordingly, the aim of this study was to investigate the effect of sleep restriction on glucose tolerance, and to examine the underlying physiological alterations that might contribute to these changes; specifically, by examining changes in mitochondrial content and function, and circadian rhythmicity. Furthermore, we examined the role of exercise as an intervention to mitigate the detrimental effects of sleep restriction. We hypothesised that sleep restriction would reduce mitochondrial content and respiratory function, and disrupt circadian rhythms, with a concomitant reduction in glucose tolerance, but that performance of HIIE would ameliorate these effects.

## Results

### Sleep Data

To verify the efficacy of our sleep interventions, we measured the participants’ total sleep time (TST) via actigraphy (33). Mean nightly TST during the intervention was significantly lower for the SR (sleep restriction) and SR+EX (sleep restriction and exercise) groups compared to the NS (normal sleep) group (*P*<0.05) (see Table 1). There was no difference in nightly TST between the SR and SR+EX group. Polysomnography (PSG), considered the gold standard assessment of sleep (34), was used to confirm actigraphy TST data and to also assess sleep architecture on night 6 of the study (n=4 per group) (Supplementary Figure 1). Both the SR and SR+EX groups obtained significantly less time in rapid eye movement (REM) sleep, non-rapid eye movement (NREM) stage 1 sleep, and NREM stage 2 sleep, compared to the NS group. Despite differences in TST between NS and both the SR and SR+EX groups, there were no significant differences in the absolute amount of sleep in the NREM stage 3 (N3) sleep between any of the groups (N3 sleep ± SD, NS = 72 ± 17, SR = 75 ± 18 min, SR+EX = 71 ± 14 min, *P* > 0.05) (Supplementary Table 1). Thus, as reported previously (2), N3 sleep was preserved despite the reduced total sleep time.

**Table 1.** Actigraphy sleep analysis and step counts for each group.

|  | NS | SR | SR+EX |
| --- | --- | --- | --- |
| **Sleep duration baseline (min)** | 448 ± 25 | 452 ± 17 | 459 ± 9 |
| **Sleep duration intervention (min)** | 449 ± 22 | 230 ± 5*<sup>#</sup> | 235 ± 5*<sup>#</sup> |
| **Step count (habitual)** | 12260 ± 3964 | 10965 ± 2136 | 11831 ± 919 |
| **Step count (intervention)** | 10652 ± 2476 | 10033 ± 1839 | 10953 ± 2316 |

Values are mean ± SD, Normal Sleep (NS), Sleep Restriction (SR) and Sleep Restriction + Exercise (SR+EX), baseline (mean of first two nights of the study), intervention (mean of 5 nights/days). \* $P < 0.05$ compared to baseline within group, # $P < 0.05$ compared to control (NS) during the intervention period.

**Figure 1.**
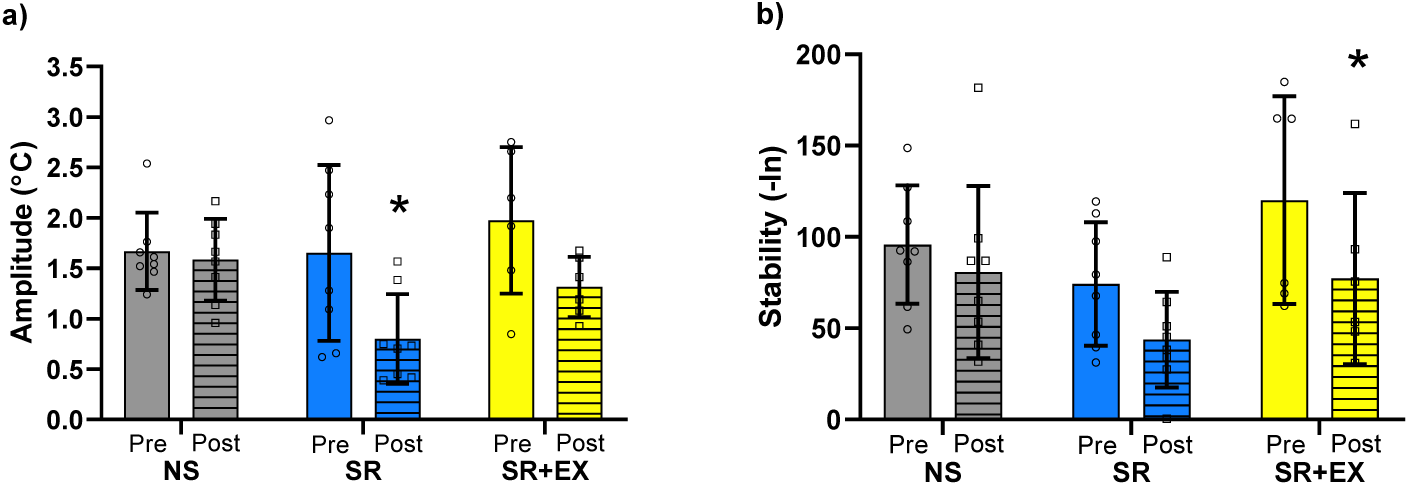
Measure of circadian peripheral skin temperature a) amplitude and b) stability pre- and post-intervention. Pre-intervention measurements are Day 2 and 3 (until 23:00 h) and the post-intervention measurements are Day 6 and Day 7. Normal Sleep (NS, *n*=8), Sleep Restriction (SR, *n*=8) and Sleep Restriction and Exercise (SR+EX, *n*=6). *Denotes significant within-group differences from pre- to post-intervention (*P*<0.05).

### Wrist skin temperature analysis

Next we used peripheral wrist skin temperature, obtained over two 48-h periods, pre- and post-intervention, to assess whether the significant reduction in TST in the SR and SR+EX groups altered aspects of circadian rhythmicity, as previously suggested (7, 8). There was a significant effect of time (*P=*0.002) for wrist skin temperature amplitude; the skin temperature amplitude in the SR group was significantly lower from pre- to post-intervention (mean amplitude change ± SD °C, 95% CI °C, *P* value; 0.85 ± 0.72°C, CI [0.20, 1.50 °C], *P*=0.008), indicating a decreased robustness of the temperature circadian rhythm. There was no change in skin temperature amplitude in the NS (0.08 ± 0.67°C, CI [-0.57, 0.73°C], *P*=0.982) or SR+EX (0.66 ± 0.65°C, CI [-0.08, 1.43°C], *P*=0.091) groups (Figure 1). Furthermore, there was a significant effect of time (*P*<0.001) for wrist skin temperature stability, with a significant reduction in the SR+EX group (53.2 ± 48.0 -ln, CI [13.7, 92.7 -ln], *P*=0.006), but not the NS (15.1 ± 39.3 -ln, CI [-21.9, 52.039 -ln], *P*=0.658) or SR (30.5 ± 30.9 -ln, CI [-6.4, 67.5 -ln], *P*=0.126) groups from pre-to post-intervention.

### Glucose tolerance

As both sleep loss and disturbances to circadian rhythm have been associated with impaired glucose tolerance, we examined plasma glucose and insulin concentrations in response to an oral glucose tolerance test (OGTT), performed before and after the sleep interventions. There were no significant differences between groups for pre-intervention glucose (*P*=0.771) and insulin (*P*=0.137) area under the curve (AUC) values. Post-intervention, there was a significant increase for total glucose AUC in the SR group (mean change ± SD, 95% CI, *P* value; 149 ± 115 A.U., CI [54, 243 A.U.], *P*=0.002) but not the NS group (−59 ± 122 A.U., CI [-36, 154 A.U.], *P*=0.356) or the SR+EX group (67 ± 57, CI [-162 to 28 A.U.], *P*=0.239). Although there was no interaction for changes in insulin AUC for any group pre-to post-intervention (*P*=0.085), there was a 29% increase in insulin AUC following the SR intervention (NS: −581 ± 1797 A.U., CI [-2037, 874 A.U.], *P*=0.933; SR: 1275 ± 1787 A.U., CI [-180, 2731 A.U.], *P*=0.100 and SR+EX: 518 ± 1043 A.U., CI [-937, 1975 A.U.], *P*>0.999) (Figure 2 and Supplementary Tables 2 and 3).

**Table 2.** Glucose, circadian, and mitochondrial-related skeletal muscle mRNA responses.

| Name | NS | SR | SR+EX | Interaction effect | Time effect |
| --- | --- | --- | --- | --- | --- |
| ***Glucose metabolism related mRNA*** |  |  |  |  |  |
| *Glut4* | -14 ± 18 | **-38 ± 33\*** | 16 ± 83 | *P*=0.192 | ***P*=0.022** |
| *Ampk1α* | -17 ± 30 | -1 ± 52 | -15 ± 24 | *P*=0.952 | *P*=0.060 |
| *Pdk4* | 0 ± 105 | 124 ± 286 | 60 ± 111 | *P*=0.862 | *P*=0.850 |
| *β-Had* | -30 ± 42 | -25 ± 39 | -15 ± 42 | *P*=0.872 | ***P*=0.011** |
| ***Circadian related mRNA*** |  |  |  |  |  |
| *Bmal1* | 2 ± 22 | **-29 ± 33\*** | 7 ± 51 | *P*=0.154 | ***P*=0.041** |
| *Clock* | 6 ± 75 | -7 ± 45 | -14 ± 37 | *P*=0.775 | *P*=0.407 |
| *Per1* | -7 ± 44 | -28 ± 48 | 2 ± 71 | *P*=0.527 | *P*=0.123 |
| *Per2* | -4 ± 32 | -7 ± 51 | -6 ± 48 | *P*=0.719 | *P*=0.410 |
| *Cry1* | 7 ± 45 | 12 ± 34 | -10 ± 21 | *P*=0.466 | *P*=0.982 |
| *Rev-Erb α* | -9 ± 38 | -9 ± 47 | -20 ± 42 | *P*=0.583 | *P*=0.130 |
| ***Mitochondrial related mRNA*** |  |  |  |  |  |
| *Pgc1α total* | -33 ± 51 | -23 ± 49 | 14 ± 61 | *P*=0.552 | *P*=0.076 |
| *p53* | 48 ± 58 | 89 ± 100 | 6 ± 41 | *P*=0.523 | *P*=0.067 |
| *Tfam* | -20 ± 33 | -22 ± 41 | -7 ± 59 | *P*=0.816 | ***P*=0.035** |
| *Nrf2* | -1 ± 34 | -11 ± 33 | -4 ± 43 | *P*=0.777 | *P*=0.179 |
| *Dnm1l* | -14 ± 45 | -3 ± 65 | -4 ± 72 | *P*=0.937 | *P*=0.170 |
| *Mfn1* | 6 ± 39 | -12 ± 47 | -5 ± 53 | *P*=0.414 | *P*=0.524 |
| *Mfn2* | 18 ± 28 | **-35 ± 28\*** | 2 ± 68 | *P*=0.562 | ***P*=0.002** |
| *Lc3b* | 4 ± 50 | -16 ± 40 | -11 ± 55 | *P*=0.486 | *P*=0.177 |
| *p62/SQSTM1* | 39 ± 75 | -1 ± 39 | -8 ± 69 | *P*=0.088 | *P*=0.628 |
| *Pink1* | -21 ± 34 | -32 ± 45 | 2 ± 74 | *P*=0.895 | *P*=0.204 |
| *Cox4* | -11 ± 22 | -23 ± 28 | -3 ± 65 | *P*=0.537 | *P*=0.088 |

Values are mean ± SD percent (%) changes from pre-intervention mRNA values. Normal Sleep (NS, *n*=7), Sleep Restriction (SR, *n*=8), and Sleep Restriction and Exercise (SR+EX, *n*=8). \*Significantly different from pre-intervention value (*P*<0.05).

**Table 3.** Baseline characteristics of participants

|  | NS<br>(n=8) | SR<br>(n=8) | SR+EX<br>(n=8) |
| --- | --- | --- | --- |
| Age (y) | 24 ± 4 | 25 ± 5 | 24 ± 4 |
| Height (cm) | 177 ± 8 | 179 ± 6 | 179 ± 7 |
| Mass (kg) | 78.7 ± 13.3 | 74.5 ± 11.7 | 80.2 ± 9.5 |
| BMI | 25.2 ± 3.6 | 23.3 ± 3.0 | 24.6 ± 2.5 |
| $\dot{V}O_{2peak}$ (mL.kg<sup>-1</sup>.min<sup>-1</sup>) | 43.7 ± 9.7 | 47.2 ± 6.7 | 48.0 ± 5.0 |
| $\dot{W}_{peak}$ (W) | 319 ± 59 | 330 ± 44 | 362 ± 48 |
| Habitual sleep duration (h:min) | 7:37 ± 0:45 | 7:08 ± 0:44 | 7:17 ± 0:39 |
| Mitochondrial respiration<br>(ETF+CI+CII)p (pmol.s<sup>-1</sup>.mg tissue<sup>-1</sup>) | 80.8 ± 17.7 | 88.3 ± 25.6 | 81.2 ± 19.7 |

Values are mean ± SD. There were no significant differences between the three groups for any of the baseline characteristics. NS – Normal sleep, SR – Sleep restriction, SR+EX – Sleep Restriction and Exercise, BMI – body mass index, $\dot{W}_{peak}$ – peak power (W), (ETF+CI+CII)p – maximal oxidative phosphorylation through electron transferring flavoprotein, complex 1 and complex 2 of the electron transport system (pre-study muscle biopsy).

**Figure 2.**
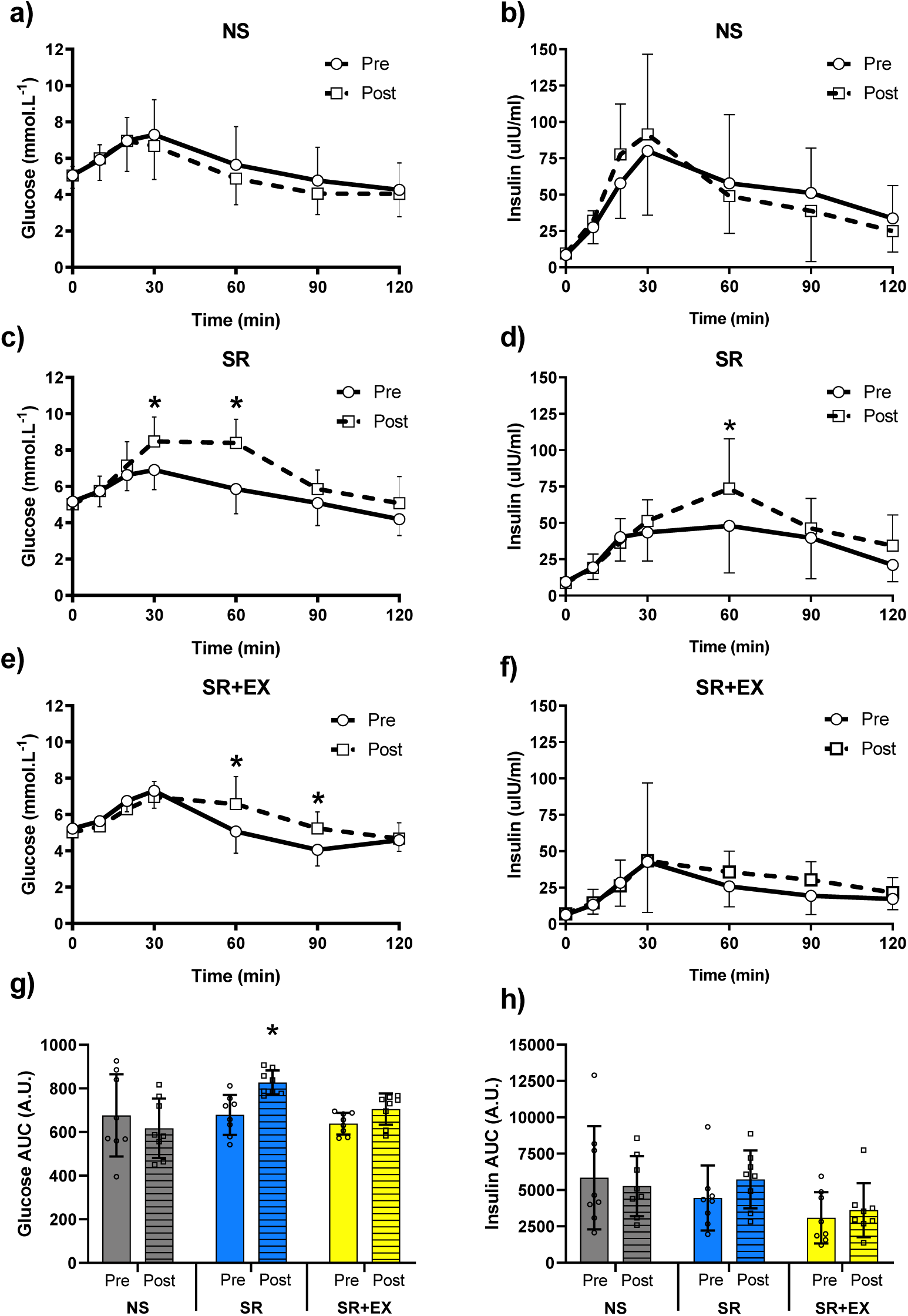
Plasma glucose and insulin concentrations for pre- and post-intervention oral glucose tolerance tests (OGTT). Plasma glucose and insulin concentrations throughout the 120-minute OGTT in the (a, b) Normal Sleep (NS), (c, d) Sleep Restriction (SR), and (e, f) Sleep Restriction and Exercise (SR+EX) groups. g) Glucose and h) Insulin total area under the curve (AUC) during the OGTT. Values are mean ± SD, individual data points are shown, * Denotes significant within-group differences from pre-to post-intervention (*P*<0.05). *n*=8 per group.

### Mitochondrial content, function, and protein synthesis

As both insulin resistance and reduced glucose tolerance have been linked to changes in mitochondrial characteristics, we investigated if skeletal muscle mitochondrial respiratory function, content, and protein synthesis (via sarcoplasmic protein synthesis) were influenced by the sleep loss and exercise interventions. There was a significant interaction effect for maximal coupled mitochondrial respiration (ETF+CI+CII)_P_ (*P*=0.032), which revealed a reduction from pre-to post-intervention (mean change ± SD pmol O_2_.s^-1^.mg^-1^, 95% CI, *P* value) in the SR group (−15.9 ± 12.4 pmol O_2_.s^-1^.mg^-1^, CI [-25.6, −6.1 pmol O_2_.s^-1^.mg^-1^], *P*=0.001) (∼18% decrease and a coefficient of variation (CV) of ∼12%) (Figure 3a). This was not evident in the NS (8.1± 6.9 pmol O_2_.s^-1^.mg^-1^, CI [-1.6 to 17.9 pmol O_2_.s^-1^.mg^-1^], *P*=0.122) or SR+EX groups (0.6 ± 11.8 pmol O_2_.s^-1^.mg^-1^, CI [-9.1, 10.4 pmol O_2_.s^-1^.mg^-1^], *P*=0.997) (Figure 3b and 3c). These results show for the first time that sleep loss is associated with decreased mitochondrial respiratory function, which was mitigated by exercise.

**Figure 3.**
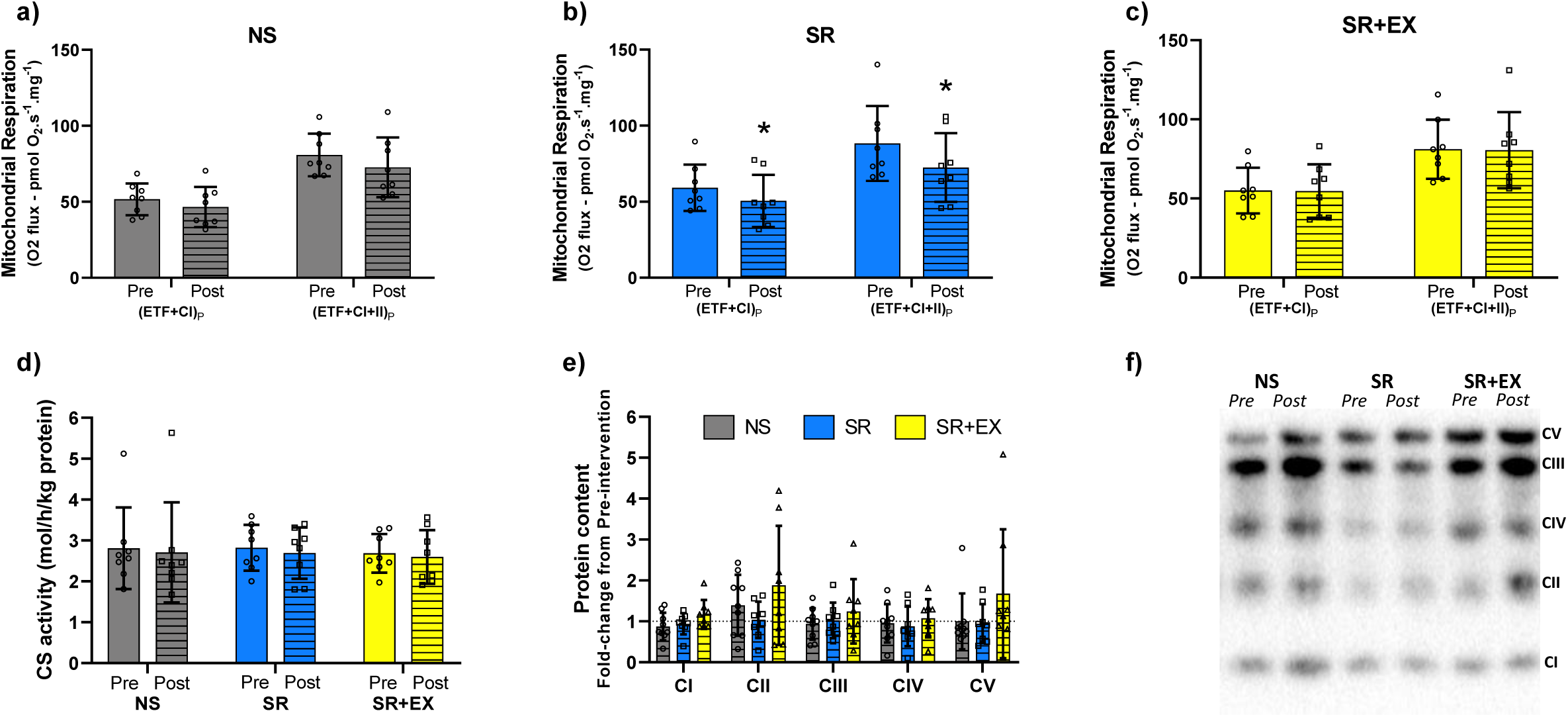
Mitochondrial respiratory function and markers of mitochondrial content from pre-intervention compared to post-intervention. Mitochondrial respiratory function in the (a) Normal Sleep (NS), (b) Sleep Restriction (SR) and (c) Sleep Restriction and Exercise (SR+EX) groups. (d) Citrate synthase activity and (e) fold-change of protein content for subunits of mitochondrial complexes (I – V) from pre-to post-intervention. Mitochondrial complex 1 (CI) - NDUFB8, Complex 2 (CII) - SDHB, Complex 3 (CIII) - Core protein 2 (UQCCRC2), Complex 4 (CIV) – MTCO, Complex 5 (CV) – ATP5A. (f) Representative image of protein content for mitochondrial complexes. Normal Sleep (NS), Sleep Restriction (SR) and Sleep Restriction + Exercise (SR+EX), (ETF+CI)_P_ – maximal coupled mitochondrial respiration through electron transfer flavoprotein (ETF) and CI; (ETF+CI+CII)_P_-maximal coupled mitochondrial respiration through ETF,CI and CII; Pre – pre-intervention, Post – post-intervention. *n*=8 per group. * Denotes significant difference within group from pre-to post-intervention (p<0.05).

Mitochondrial content can be assessed via CS activity and the protein content of mitochondrial complex subunits (16). There were no changes in whole-muscle CS activity from pre-to post-intervention for any of the groups (interaction, *P*=0.972) (mean difference, 95% CI, *P* value, NS 0.10 ± 0.45 mol/h/kg protein, CI [-0.24, 0.44 mol/h/kg protein], *P*>0.999); (SR 0.13 ± 0.29 mol/h/kg protein, CI [-0.21, 0.47 mol/h/kg protein], *P*=0.992) and (SR+EX 0.08, ± 0.35 mol/h/kg protein, CI [-0.26, 0.43 mol/h/kg protein], *P*>0.999) (Figure 3d). As a further validation of changes in mitochondrial content, the protein content for subunits of mitochondrial complexes were assessed via western blotting. No significant interaction effects were observed for the protein content of Complex 1 (*P*=0.116), Complex 2 (*P*=0.649), Complex 3 (*P*=0.621), Complex 4 (*P*=0.718) or Complex 5 (*P*=0.158) (Figure 3e), collectively indicating that the sleep and exercise interventions did not affect mitochondrial content.

It has been argued that the best measure of mitochondrial biogenesis is mitochondrial protein synthesis (MitoPS) (35). The sarcoplasm (skeletal muscle cytoplasm) is enriched with mitochondria (see Figure 4a), and protein synthesis within this fraction (sarcoplasmic protein synthesis - SarcPS) is likely reflective of MitoPS (24). Therefore we assessed, for the first time, the effects of sleep restriction on SarcPS, which was significantly lower in the SR group compared to both the NS group (between groups difference FSR %/day ± SD, 95% CI, *P* value, −0.62 ± 0.11%, CI [−0.90, -0.33], *P*<0.001) and the SR+EX group (−0.66 ± 0.12%, CI [−0.95, -0.37], *P*<0.001); there was no difference in SarcPS between the NS and SR+EX groups (0.04 ± 0.10%, CI [-0.33, 0.24], *P*>0.999) (Figure 4b). These new data suggest that exercise can mitigate the lower rates of SarcPS seen following sleep restriction alone.

**Figure 4.**
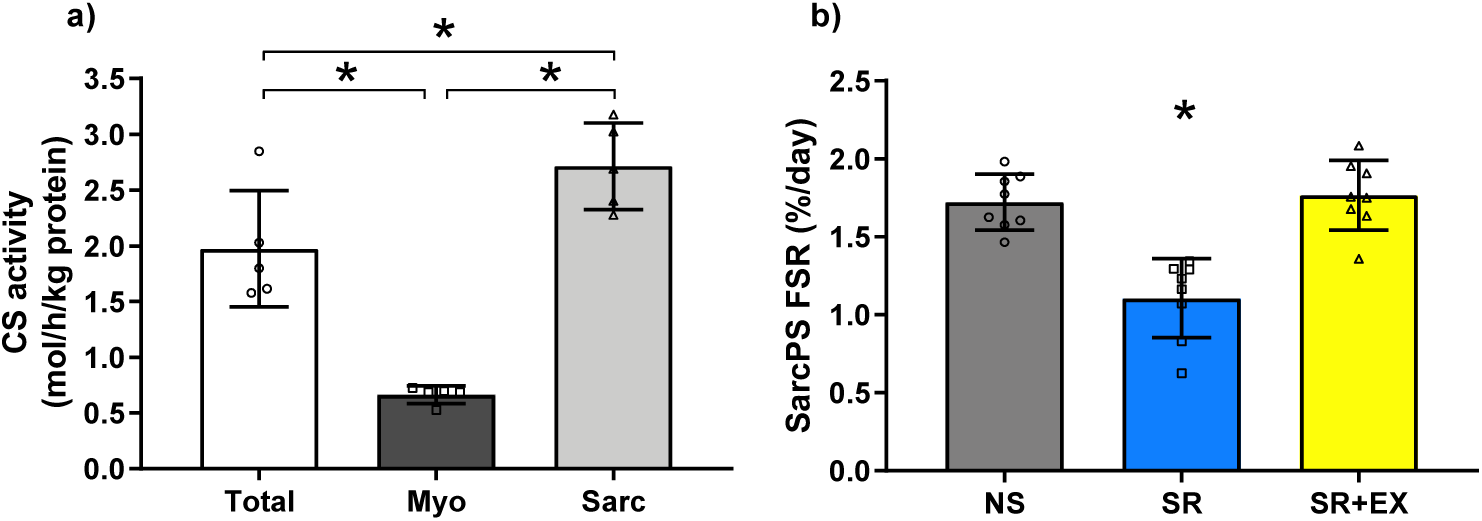
Citrate synthase (CS) activity from fractionated skeletal muscle samples and Sarcoplasmic protein synthesis (SarcPS) –. a) CS activity of whole-muscle lysate (Total), myofibrillar (Myo), and sarcoplasmic (Sarc) fractions were assessed from the same muscle samples (*n*=5). b) Fractional synthetic rate (FSR) of SarcPS during the sleep intervention. Data are mean ± SD. Normal Sleep (NS), Sleep Restriction (SR), and Sleep Restriction + Exercise (SR+EX), *n*=8 per group. *denotes significantly different from other groups (*P*<0.05).

### Glucose, circadian, and mitochondrial-related gene expression

Given the changes observed for mitochondrial respiration, protein synthesis, circadian rhythm (reflected by peripheral skin temperature), and glucose tolerance, we investigated the expression of genes that regulate these processes (Table 2). In the SR group, there was a significant reduction from pre-to post-intervention in the mRNA of *Mfn2* (mean change ± SD %, 95 % CI, *P* value; −35 ± 28 %, CI [-7, 79 %], *P*=0.044), *Bmal1* (−29 ± 33%, CI [-6, 64%] *P*=0.031), and *Glut4* (−38 ± 33%, CI [-13, 88%], *P*=0.020), which were not evident in the NS or SR+EX groups. There was also a decrease in *Tfam* and *β-Had* mRNA expression with time; however, post-hoc analysis revealed no specific group differences from pre-to post-intervention.

### Glucose, circadian, and mitochondrial-related protein content

Following our discovery of reductions in the mRNA of some mitochondrial, circadian, and glucose-related genes, we used the limited amount of remaining muscle biopsy tissue to assess whether this leads to further changes at the protein level. There were no significant interaction effects for PGC-1α (*P*=0.257), DRP1 (*P*=0.642), MFN2 (*P*=0.768), p53 (*P*=0.294), BMAL1 (*P*=0.778), or GLUT4 (*P*=0.466) protein content, from pre-to post-intervention (Figure 5).

**Figure 5.**
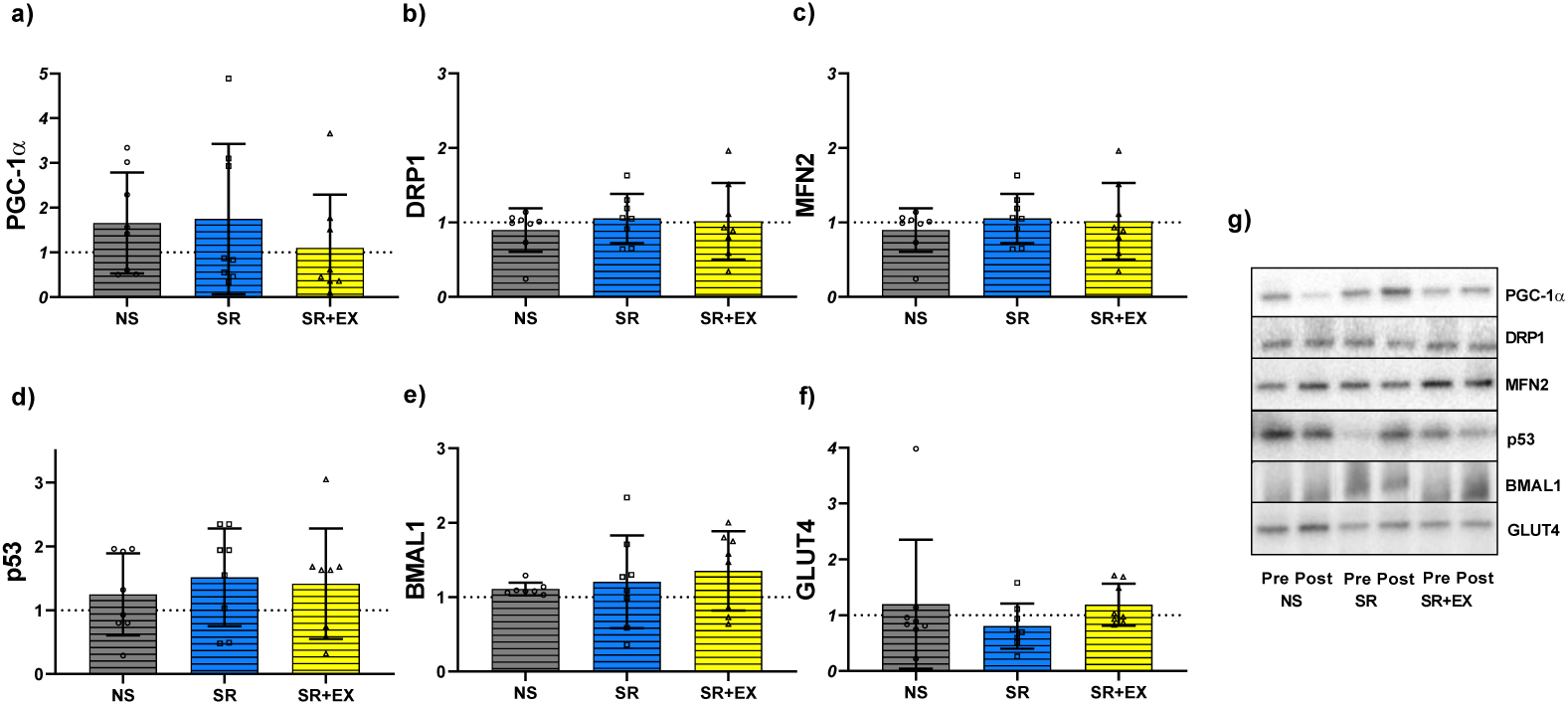
Skeletal muscle mitochondrial, circadian, and glucose-related protein content. a) PGC-1α, b) DRP1, c) MFN2, d) p53, e) BMAL1, f) GLUT4, and g) representative western blot images. Data are mean values ± SD, normalised to pre-intervention values. Normal Sleep (NS), Sleep Restriction (SR) and Sleep Restriction and Exercise (SR+EX), *n*=8 per group. * Denotes significant change from pre-intervention (*P*<0.05).

## Discussion

We discovered that in healthy young men, sleep restriction resulted in significantly impaired glucose tolerance, with concomitant changes in circadian rhythmicity (skin temperature amplitude and clock gene expression), skeletal muscle mitochondrial respiratory function, and sarcoplasmic protein synthesis (SarcPS) – a proxy for mitochondrial protein synthesis. However, performing three sessions of high-intensity interval exercise (HIIE) during the sleep restriction intervention mitigated the perturbations that were observed in the SR group. Mechanistically, there were also differences between the SR and SR+EX groups in the expression and content of essential glucose and mitochondrial-related regulatory markers. Our study provides novel insights into the potential mechanisms underlying previously reported changes in glucose tolerance with sleep loss and suggests exercise may be used as a therapeutic intervention to attenuate such adverse effects.

The sleep restriction protocol used in this study (consisting of 4 h TIB per night, for five consecutive nights) resulted in a significant impairment in glucose tolerance. Furthermore, the plasma insulin response during the OGTT for the SR group increased by 29% from pre-to post-intervention. These results are consistent with previous studies (3-5), including Rao et al. (2) who reported a 25% decrease in whole-body insulin sensitivity (measured with a hyperinsulinaemic-euglycaemic clamp) following a similar sleep restriction protocol (2). Therefore, the 22% increase in plasma glucose AUC following sleep restriction observed in young, healthy men in this study solidifies the evidence supporting the detrimental effect of even short periods of sleep restriction on glucose tolerance.

A novel aspect of this study was to examine the effect of three sessions of HIIE during the period of sleep restriction as a means of mitigating sleep restriction-induced reductions in glucose tolerance. In contrast to the SR group, adverse changes in glucose tolerance were mitigated in the SR+EX group. While others have shown acute positive (36) and protective (37) effects of exercise on sleep-loss-induced changes in glucose tolerance, these studies assessed glucose tolerance either immediately after or within 24 h of exercise, raising the possibility these findings were confounded by the acute effects of exercise on glycaemic control, which are known to persist for up to 48 h (22). In contrast, we performed the OGTT 48 h post exercise, thus demonstrating for the first time that performing HIIE during a period of sleep restriction prevents the negative effects to glucose tolerance.

The changes in glucose tolerance were mirrored by changes in skeletal muscle glucose transporter 4 (*Glut4*) mRNA expression, whereby *Glut4* was lower in the SR group but maintained in the SR+EX group. The maintenance of *Glut4* expression in the SR+EX group is likely explained by the well-documented increase in expression commonly observed with exercise (38). While there were no changes in GLUT4 protein content in our study, a previous study that used streptozocin-induced diabetic rats demonstrates that changes in skeletal muscle *Glut4* mRNA coincide with reductions in *in-vivo* glucose uptake, but precede changes in GLUT4 protein content (39). Given that there was no change in GLUT4 protein content in these previous studies, it may suggest that GLUT4 translocation may be impaired and supports previous research indicating impairments within the insulin signalling pathway, following sleep restriction (40, 41).

Insulin resistance and reductions in glucose tolerance have previously been linked to altered mitochondrial characteristics (14, 15); therefore, we also assessed changes in skeletal muscle mitochondrial respiratory function. To our knowledge, ours is the first report of a SR-induced reduction in skeletal muscle mitochondrial respiratory function in humans. Previously, a study in humans investigating a single night of 4 h TIB reported a decrease in insulin sensitivity with a concomitant increase in plasma acylcarnitines (9), which was suggested to be indicative of both reduced fatty acid oxidation and impaired mitochondrial function. The maintenance of mitochondrial respiratory function in the SR+EX group is consistent with previous reports of the potency of HIIE for improving mitochondrial respiratory function (26, 42); however, we admittedly do not have a Normal Sleep and HIIE group that would allow us to ascertain whether HIIE enhanced mitochondrial function. Nonetheless, our results do provide evidence supportive of a link between reduced mitochondrial respiratory function and impaired glucose tolerance with sleep restriction that is mitigated by the addition of HIIE.

Mitochondria are dynamic organelles and the production of new mitochondrial proteins is an important determinant of their overall function (35). Mitochondrial protein synthesis (MitoPS) is the measure that best reflects the process of mitochondrial biogenesis (23, 35). However, tissue availability for this project necessitated the use of sarcoplasmic protein synthetic rate (e.g., SarcPS – Figure 4a) as a proxy for MitoPS. For the first time, we report a lower rate of SarcPS in the SR group compared to both the NS and SR+EX groups (Figure 4b). The significance of the lower rate of SarcPS in the SR group is difficult to ascertain; however, we hypothesise it may underpin a reduced turnover of mitochondrial proteins, with a subsequent impact on mitochondrial respiratory function (35). Support for this notion comes from the observation that changes in MitoPS and mitochondrial respiratory function have been shown to occur concomitantly (23, 43); nonetheless, a direct link is yet to be established. Moreover, HIIE has consistently been reported to increase both SarcPS and MitoPS (24, 44), and this likely accounts for the higher rate of SarcPS in the SR+EX group compared to the SR group. Indeed, previous reports indicate that SarcPS remains elevated for 48 h following endurance exercise (45), and these increases are likely reflected in the integrative measure of protein synthesis used in this study, which represents a summation of SarcPS throughout the entire intervention.

Considering the changes observed to SarcPS, the effect of sleep restriction on markers of mitochondrial content was also assessed. There were no changes to the protein content of mitochondrial complex subunits or CS activity – both of which are valid markers of mitochondrial content (16). While reduced CS activity has previously been reported following sleep interventions, this was in response to extreme sleep deprivation (i.e., 120 h of continuous wakefulness) (18); thus, a direct comparison with our findings is difficult. Despite no change in CS activity or protein content of mitochondrial complex subunits in the SR+EX group, endurance exercise is well known to increase mitochondrial content (25). Our results suggest changes in mitochondrial respiratory function and SarcPS occurred independently of detectable changes in mitochondrial content. This dissociation between changes in mitochondrial content and respiratory function has previously been reported, and it has been suggested that these properties may be differentially regulated (26, 43). One explanation for our observations may involve the processes regulating mitochondrial dynamics/remodelling (i.e., fission and fusion), which can alter the efficiency and oxidative capacity of the mitochondria, without necessarily altering mitochondrial content (23). Therefore, we assessed genes and proteins known to regulate mitochondrial remodelling. The reduced Mitofusin 2 (*Mfn2*) gene expression in this study points to a potential link between altered mitochondrial morphology and the reduced mitochondrial respiratory function and glucose tolerance observed in the SR group. Considering exercise can also influence mitochondrial dynamics and mitophagy (46, 47), and that *Mfn2* has previously been shown to be elevated in skeletal muscle 24 h post an endurance exercise session (48), this may explain why *Mfn2* mRNA levels were maintained in the SR+EX group.

As many of the molecular processes that regulate mitochondrial content, function, and dynamics are regulated in a circadian manner (49, 50), we also examined aspects of circadian rhythmicity using the robust measures of peripheral skin temperature (obtained over 48 h, pre- and post-intervention) and analysis of molecular clock gene expression. Peripheral skin temperature measures oscillate inversely to the rhythms of core body temperature and therefore provides a physiological circadian output. We show that skin temperature amplitude, a common measure of assessing the circadian nature of biological processes, was significantly reduced in the SR group. Our data parallels that of Moller-Levet et al. (51) who reported in human white blood cells, reduced amplitude of core clock gene circadian expression following seven nights of sleep restriction (6 h TIB each night). At the skeletal muscle level, we also show a significant reduction in *Bmal1* mRNA from pre-to post-intervention in the SR group, but not in the NS or SR+EX groups. This supports previous findings from Cedernaes et al. (7, 8) who reported a decrease in *Bmal1* mRNA expression following 24 h of sleep deprivation. Moreover, reductions in *Glut4* mRNA expression, similar to those we report in the SR group, have also been reported in *Bmal1*^-/-^ mice (52), suggesting the reductions in *Bmal1* mRNA and changes to circadian rhythm that were observed in the SR group may contribute to changes in *Glut4* mRNA. These findings suggest that sleep restriction can disrupt the skeletal muscle circadian clock, which has been associated with negative metabolic consequences, such as insulin resistance and mitochondrial function (21, 52).

Exercise is considered a potent zeitgeber (i.e., circadian time cue) capable of altering circadian rhythms (31, 53), therefore, we examined its effect on skin temperature and clock gene expression. In this study, there was a reduction in skin temperature stability, whilst skeletal muscle *Bmal1* mRNA expression was maintained in the SR+EX group, but not the SR group. Exercise-like contractile activity in C2C12 myotubes induce time of day related-phase shifts in *Bmal1* mRNA expression (53), which may help to explain our findings. Nevertheless, further studies and additional muscle sampling time-points are needed to clarify the effect of exercise performed at different times of the day in humans, as well as changes in the underlying regulatory mechanisms (i.e., clock gene expression). This will further optimise the implementation of exercise strategies to realign changes in circadian rhythms induced by inadequate sleep.

In summary, we have provided the first direct evidence of a concomitant decrease in mitochondrial respiratory function, SarcPS (of which MitoPS is a contributor), and glucose tolerance, following sleep restriction in otherwise healthy young men. We discovered alterations in mRNA expression of select genes involved in glucose uptake, mitochondrial dynamics, and circadian rhythms (Figure 6). Collectively, these results highlight a number of potential mechanisms by which sleep restriction may lead to reductions in glucose tolerance. Importantly, HIIE mitigated these detrimental changes in glucose tolerance and mitochondrial characteristics. While further research is still required, these data provide a basis for the development of evidence-based health guidelines and recommendations for those experiencing inadequate sleep, by highlighting some of the underlying biological mechanisms that can be targeted by therapeutic interventions such as exercise.

**Figure 6.**
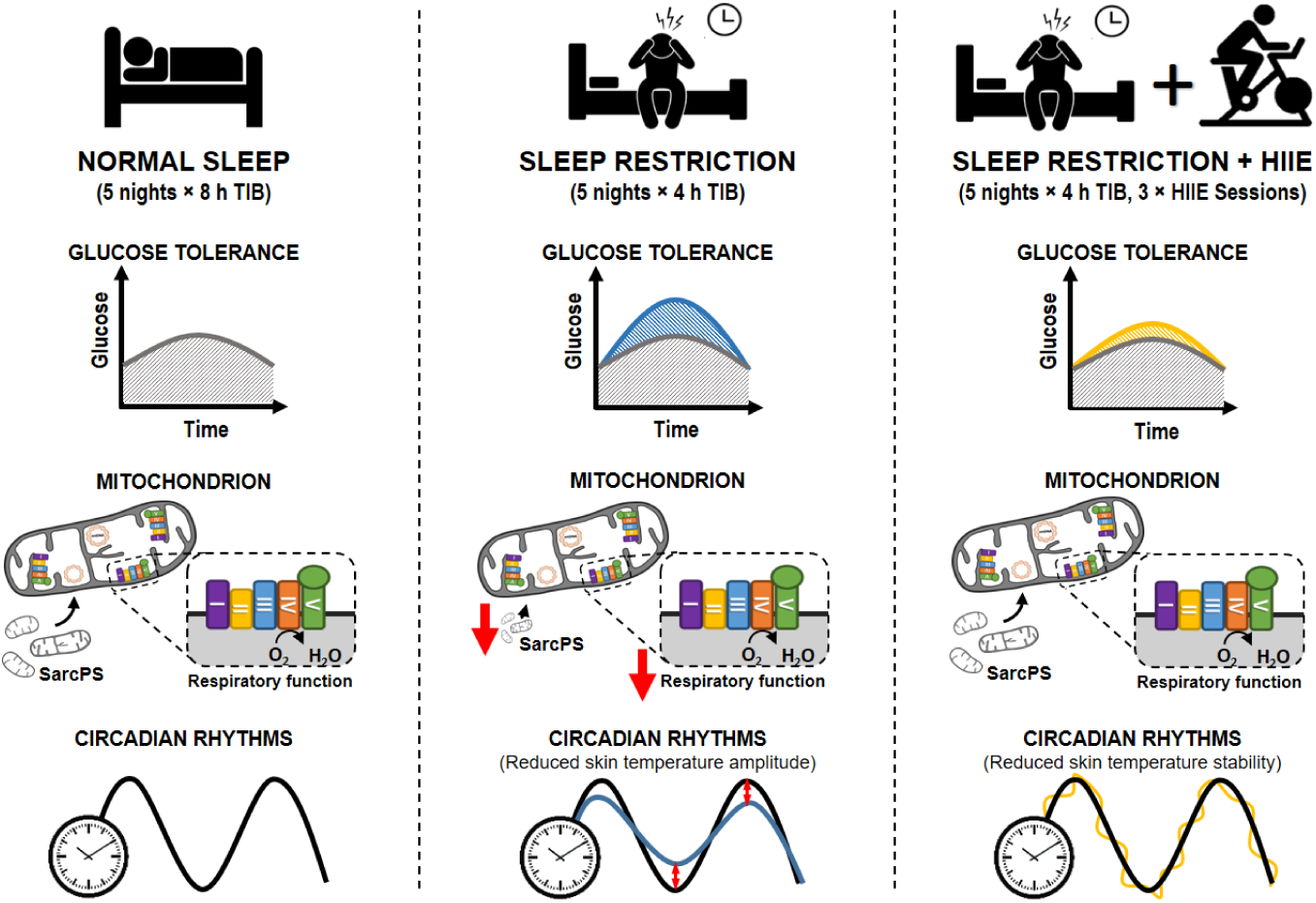
Summary of the effects of sleep restriction, with or without HIIE, on glucose tolerance, mitochondrial characteristics, and circadian rhythms. NS – Normal Sleep group, SR – Sleep Restriction group, SR+EX – Sleep Restriction and Exercise group, HIIE – High-intensity interval exercise, SarcPS – Sarcoplasmic protein synthesis, TIB – Time in bed.

## Study methodology

### Ethics approval

All procedures involved conform to the standards set by the latest revision of the Declaration of Helsinki (except for registration in a database) and were approved by the Victoria University Human Research Ethics Committee (HRE15-294).

### Participants

Twenty-four healthy, recreationally active men, aged between 18 and 40 years of age, volunteered to participate. Eligible participants 1) were not taking any medications, 2) were not performing shift-work (within the previous three months), 3) had regular sleeping habits (6 – 9 hours per night) and no previously diagnosed sleep disorders, 4) had not travelled overseas in the previous two months, and 5) had a body mass index between 19 and 30.

### Study overview

Eligible participants attended the exercise physiology laboratory for baseline anthropometric measurements (i.e., height and body mass), and aerobic fitness testing (peak oxygen uptake 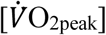 and peak aerobic power [Ẇ_Peak_]) that was performed to volitional exhaustion on an electronically braked cycle ergometer (Excalibur, V2.0; Lode, Groningen, Netherlands), using an incremental ramp protocol (30 W/minute). One week prior to the intervention, a resting skeletal muscle biopsy was obtained to determine baseline levels of deuterium oxide (D_2_O) enrichment, and to assess basal mitochondrial respiratory function. Following baseline testing, participants were matched for age, BMI, habitual sleep duration, 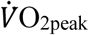, and mitochondrial respiratory function, and then allocated to one of three experimental groups, in a counterbalanced order (Table 3).

The study consisted of an eight-night stay within a temperature-controlled sleep laboratory. All groups completed two initial nights of baseline sleep (8 h TIB from 23:00 h to 07:00 h), followed by a five-night intervention period, during which the NS group spent 8 h TIB (23:00 h to 07:00 h), while both SR and SR+EX spent 4 h TIB per night (03:00 h to 07:00 h). Between 23:00 h and 03:00 h, lighting was dimmed to below 15 lux to reduce the effect of lighting on circadian rhythms (54). The SR+EX group also performed three exercise sessions during the intervention period on days 4, 5, and 6 at 10:00 h. Following the intervention period, all groups completed a final night of *ad libitum* recovery sleep. Participants were monitored throughout the protocol and provided with a standardised diet consisting of fixed proportions (relative to body mass) of carbohydrates (4.5 g.kg^-1.^d^-1^), protein (1.5 g.kg^-1.^d^-1^) and fat (1 g.kg^-1.^d^-1^). All mealtimes (six throughout the day) were kept constant. An overview of the study protocol is shown in Figure 7.

**Figure 7.**
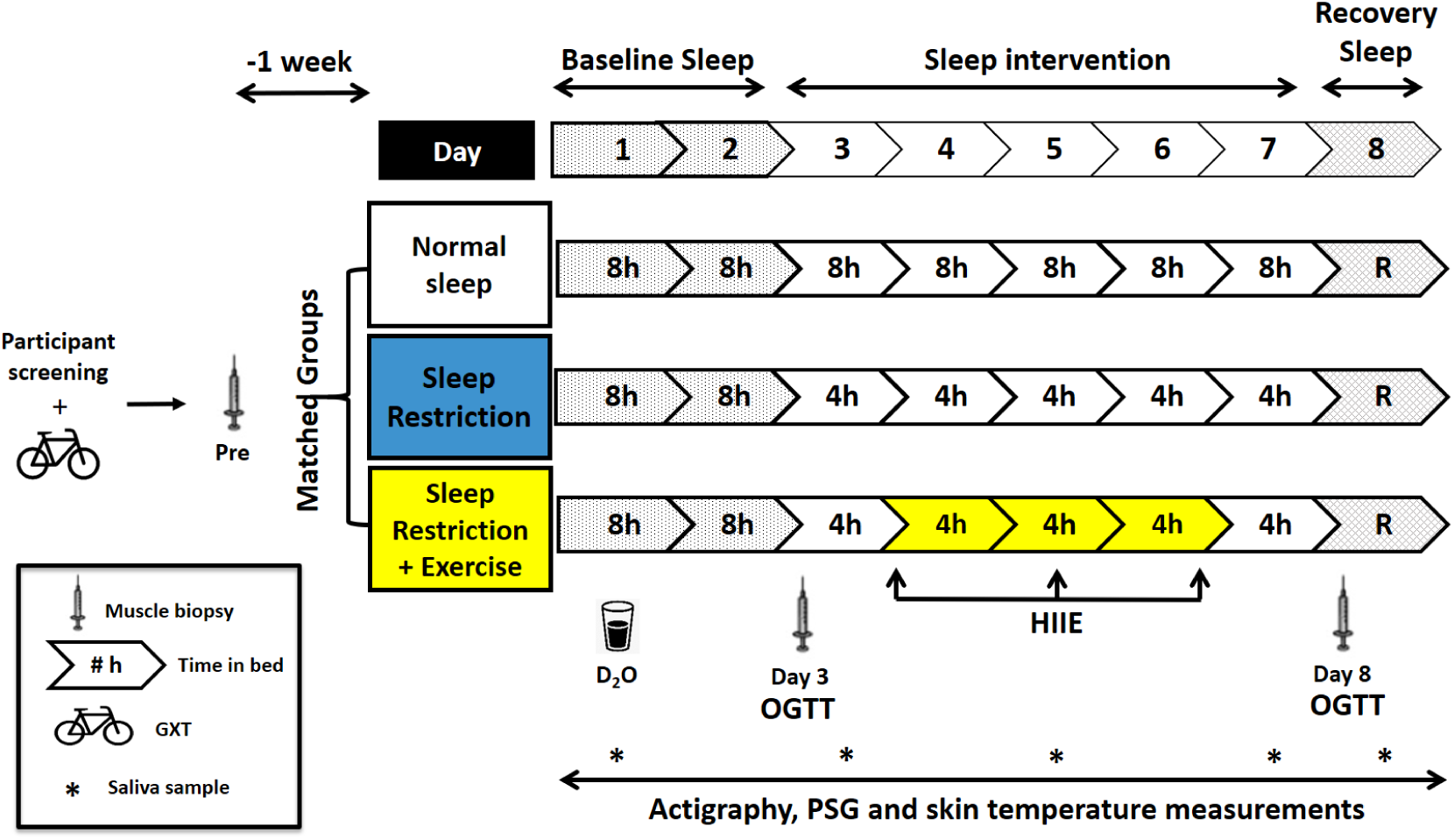
Schematic representation of the study protocol. OGTT – oral glucose tolerance test, GXT – Graded exercise test, D_2_O – deuterium oxide ingestion, HIIE – high-intensity interval exercise, R – *ad libitum* recovery sleep, PSG – polysomnography sleep monitoring, participant screening refers to medical questionnaires, exclusion criteria and, habitual sleep, and physical activity monitoring.

## Experimental procedures

### Sleep and physical activity monitoring

Sleep was assessed for one week prior to and then throughout the study using wrist-watch activity devices (Actiwatch 2, Philips Respironics, Murrysville, PA, USA) (33). Sleep architecture, duration, and quality were also determined via polysomnography (PSG) (Compumedics, AUS) on night 6 for a subset of participants from each condition (*n*=12; 4 per group). Electrode placement for PSG monitoring was determined using the 10-20 electrode placement system and scored in accordance to standard criteria (55). Habitual daily step counts were monitored using validated step-counting applications on the participants’ personal mobile phone devices (i-Health app, Apple Inc., Cupertino, CA, USA; and Samsung Health, Samsung Electronics Co., Ltd., Suwon, South Korea) and these step counts were replicated throughout the study (Table 1) (data modified from (56)).

### High-intensity interval exercise (HIIE)

The HIIE protocol was adapted from previous studies, which demonstrated improvements in glucose tolerance (27, 57) and consisted of 10 × 60-second intervals performed on a cycle ergometer (Velotron, Racer-Mate, Seattle, WA, USA) at 90% of each participant’s Ẇ_peak_. Each interval was interspersed with 75 seconds of active recovery at 60 W. Each session started with a 3-minute warm up at 60 W. The mean power per interval was 318 ± 53 W and the mean HR throughout the protocol was 156 ± 13 bpm.

### Wrist skin temperature measurements

Wrist skin temperature was measured every 10 min (at a sensitivity of 0.0625°C) across two 48 h periods (Pre-intervention - Days 2 and 3 (until 11 pm) and Post-intervention - Days 6 and 7), using non-invasive temperature recording devices (iButtons, Thermochron iButton; Embedded Data Systems, Lawrenceburg, KY). This method has been shown to be reliable and valid for evaluating temperature circadian rhythmicity, with peripheral wrist skin temperature reported to have an inverse relationship to core body temperature (58, 59). The data was analysed as previously described (59), for variations in temperature amplitude and stability for each participant, using a modified version of the software program JTK_CYCLE (60). Temperature amplitude is defined as the difference between the maximum (or minimum) value of the trace and the mesor (which represents the mean value of the data after smoothing with a cosine function) across a 24-h period (61); changes in amplitude having previously been proposed to provide an indication of the robustness of the rhythm (58, 62). The degree of phase homogeneity of a rhythm during the period of data collection is considered a description of the ‘stability’ of the rhythm (i.e., stability is considered high when the oscillatory pattern of the rhythm is nearly identical from one day to the next, and considered low when there are large discrepancies between oscillatory patterns from day-to-day) (58, 62). Data from eight participants in both the NS and SR groups were included; however, due to technical issues, data from the SR+EX group includes only six participants.

### Oral glucose tolerance testing (OGTT)

To assess glucose tolerance, OGTT tests were performed on Day 3 (pre-intervention) and Day 8 (post-intervention) at 08:00 h, following an overnight fast. Participants consumed a 300 mL solution containing 75 g of glucose (Point of Care Diagnostics-Scientific, NSW, Australia) and blood samples were collected after 0, 10, 20, 30, 60, 90, and 120 minutes. Plasma glucose concentrations were measured on a glucose/lactate analyser (YSI, 2300 STAT plus, Yellow Spring, OH, USA). Plasma insulin concentrations were assessed using an Insulin ELISA kit (ALPCO. 80-INSHU-E01.1, E10.1, Salem, NH, USA) and run according to the manufacturer’s instructions.

### Muscle biopsies

One week prior to commencing the study, and on Day 3 and Day 8 of the intervention, muscle biopsies were sampled from the *vastus lateralis* muscle using a suction-modified Bergström needle, and under local anaesthesia of the skin and fascia (1% lidocaine). All samples were collected at 10:00 h. Samples on Day 3 and 8 were collected post OGTT. All samples were immediately frozen in liquid nitrogen and stored at −80°C, or set aside for mitochondrial respirometry.

### High-resolution respirometry

Immediately following the muscle biopsies, muscle fibres were placed in ice-cold biopsy preserving solution (BioPS) and prepared as previously described (26). Mitochondrial respiration was measured in triplicate (coefficient of variation [CV] = 12%) (from 2 to 4 mg wet weight of muscle fibers) in MiR05 at 37°C by using the high-resolution Oxygraph-2k (Oroboros, Innsbruck, Austria). Oxygen concentration (in nanomoles per milliliter) and flux (in picomoles per second per milligram) were recorded with DatLab software (Oroboros). The following substrate–uncoupler–inhibitor titration (SUIT) protocol was used to assess mitochondrial respiratory function; octanoyl-carnitine (0.2 mM) and malate (2 mM) for leak respiration (L) via electron transferring flavoprotein (ETF), (ETF)L, ADP (5 mM, saturating concentration) was added for measurement of oxidative phosphorylation capacity (P), (ETF)P. Pyruvate (5 mM) was then added for measurement through Complex 1 (CI), (ETF+CI)P, followed by addition of succinate (10 mM) for measurement of oxidative phosphorylation capacity through complex 1 and 2 combined (ETF+CI+II)P. Cytochrome c (10 mM) was then added to test for outer mitochondrial membrane integrity (an oxygen flux increase of < 15% from (ETF+CI+II)P was considered acceptable). A series of stepwise carbonyl cyanide 4-(trifluoromethoxy) phenylhydrazone (FCCP) titrations (0.75–1.5 mM), for the measurement of ETS capacity (E) through (ETF+CI+II)E followed. Rotenone (0.5 mM), an inhibitor of CI, was then added to determine E through CII (ETF+CII)E, whereas addition of antimycin A (2.5 mM), an inhibitor of CIII, allowed measurement of and correction for residual oxygen consumption (ROX), which is indicative of non-mitochondrial oxygen consumption. As there was no difference in mitochondrial respiration between ‘Pre’ and Day 3 resting biopsies (mean (ETF+CI+II)P pmol O_2_.s^-1^.mg^-1^ ± SD; Pre – 81.4 ± 21.6 pmol O_2_.s^-1^.mg^-^ 1, Day 3 - 80.6 ± 21.0 pmol O_2_.s^-1^.mg^-1^, *P*=0.831), ‘Baseline’ mitochondrial respiration was calculated as the mean from the ‘Pre’ study and Day 3 muscle biopsies.

### Assessment of sarcoplasmic protein synthesis (SarcPS)

SarcPS was used as an indicator of mitochondrial protein synthesis (MitoPS), as performed previously (24), due to the large amount of muscle needed to asses MitoPS (e.g., 80 – 100 mg). In validating this approach, we measured CS activity (a validated biomarker of mitochondrial content (16)) in whole-muscle lysate, as well as the myofibrillar and sarcoplasmic fractions (Figure 4a). The whole-muscle CS activity was significantly higher than the myofibrillar fraction (mean difference CS activity (mol/h/kg protein) ± SD, 1.30 ± 0.48, CI [0.64, 1.97], *P*<0.001). The sarcoplasmic fraction CS activity was also significantly higher than the myofibrillar fraction (2.05 ± 0.35, CI [-2.71, −1.39], *P*<0.001) and whole-muscle fractions (0.74 ± 0.44, CI [−1.40, -0.07], *P*=0.027) demonstrating that the sarcoplasmic fractions are enriched with mitochondria, compared to both the myofibrillar fraction and whole-muscle sample.

To determine SarcPS, on Day 1 each participant ingested 150 mL of Deuterium Oxide (D_2_O) (70 atom %, Cambridge Isotope Laboratories) as previously described (63). Saliva samples were collected prior to D_2_O ingestion and then on Days 3, 5, 7, and 8 to determine body water enrichment via cavity ring-down spectroscopy (Picarro L2130-I analyser, Picarro, Santa Clara, CA). Total body water ^2^H enrichment was used as a surrogate for plasma alanine ^2^H labelling, as previously described (63). The mean body water enrichment (atom percent excess, APE) has been reported previously (56).

Frozen muscle samples (40 to 60 mg) were homogenised and prepared as previously described (64). Cation exchange chromatography was then performed on the sarcoplasmic samples using columns containing Dowex resin (Dowex 50wx8-200 ion exchange resin, Sigma Aldrich) to extract free amino acids from the sarcoplasmic fractions (64). The amino acid samples were then derivatised as their N-acetyl-n-propyl-esters, as per previous protocols (65).

The ^2^H/^1^H ratio of the sarcoplasmic samples were determined using gas chromatography pyrolysis isotope ratio mass spectrometry (GC-P-IMS) (Metabolic Solutions, Nashua, NH, USA), to assess the incorporation of deuterium into protein-bound alanine. This was used to assess the fractional synthetic rate (FSR) of sarcoplasmic proteins with the use of the enrichment of body water, corrected for the mean number of deuterium moieties incorporated per alanine (i.e., 3.7) as previously described (63), as the surrogate precursor labelling between subsequent biopsies.

The following standard equation (63) was used to determine FSR:

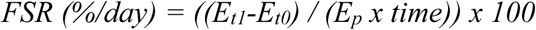

Where; FSR = fractional synthetic rate, E_t1_ = APE day 8, E_t0_ = APE day 3, E_p_ = average saliva APE, time = time between biopsies, in days, and APE = atomic percentage excess.

### Preparation of whole-muscle lysates for western blots and CS activity assay

Frozen muscle (10 to 20 mg) was homogenised as previously described (26) in an ice-cold lysis buffer (1:20 w/v) containing 50 mM Tris, 150 mM NaCl, 1 mM EDTA, 1% IGEPAL, deionised water and a protease/phosphatase inhibitor cocktail (Cell Signaling Technology (CST), Danvers, MA, USA), adjusted to pH 7.4. Protein concentration was determined in triplicate with a commercial colorimetric assay (Protein Assay kit-II; Bio-Rad, Gladesville, NSW, Australia), against bovine serum albumin standards (BSA, A9647; Sigma-Aldrich).

### Citrate synthase activity assay

Citrate synthase (CS) activity was determined in triplicate on a 96-well microtiter plate by adding 7.5 μL of a 4 mg/mL muscle homogenate (freeze thawed in liquid nitrogen twice), 40 μL of 3 mM acetyl CoA, 25 μL of 1 mM 5,59-dithiobis(2-nitrobenzoic acid) (DTNB), 165 μL of 100 mM Tris buffer (pH 8.3, kept at 30 °C). After addition of 15 μL of 10 mM oxaloacetic acid, the plate was immediately placed in an xMark-Microplate spectrophotometer (Bio-Rad) at 30°C, and after 30 s of linear agitation, absorbance at 412 nm was recorded every 15 s for 3 min. CS activity is reported as moles per hour per kilogram of protein.

### Western blotting

Muscle homogenate was diluted in 4X Laemmli buffer, and equal amounts of total protein (15 or 20 μg) were loaded in different wells on Criterion™ 4-20% TGX Stain-Free™ Precast Gels (Bio-Rad, Australia). A stain-free system (Bio-Rad, Australia) was used as a loading control, with protein expression normalised to total protein loaded per lane. Each gel also contained four to six internal standards of varying dilutions, made from a mixed homogenate of every sample in equal concentrations. These standards were used to form a calibration curve, with density plotted against protein content. Protein abundance was then calculated from the measured band intensity for each sample on the gel, using the linear regression equation from the calibration curve.

Muscle lysates were separated via gel electrophoresis and transferred to PVDF membranes. Membranes were then blocked in 5% non-fat dry milk diluted in Tris-buffered saline with 0.1% Tween-20 (TBST) for 60 minutes. Membranes were then incubated overnight at 4°C with the appropriate primary antibody, prepared at a 1:1000 dilution in TBST with 5% BSA and 0.02% sodium azide (unless stated otherwise). The following primary antibodies were from Cell Signaling Technologies (CST) and include PGC-1α (CST2178), DRP1 (CST5341), MFN2 (CST9482), p53 (CST2527), GLUT4 (CST2213). The following antibodies were obtained from Abcam; BMAL1 (AB93806), Total OXPHOS (AB110413). The membranes were then incubated at room temperature with the appropriate host species–specific secondary antibody for 60 min, before being exposed to a chemiluminescence solution (Clarity(tm) Western ECL Substrate [Bio-Rad, Hercules, CA, USA] or SuperSignal(tm) West Femto Maximum Sensitivity Substrate [ThermoFisher, ThermoFischer Scientific, Wilmington, DE, USA]). Images were taken with a ChemiDoc Imaging System fitted (Bio-Rad). Densitometry was performed with Image Lab 5.0 software (Bio-Rad). Images are typically displayed with at least five bandwidths above and below the band of interest.

### Real-time quantitative polymerase chain reaction (qPCR)

RNA extraction with TRIzol (Life Technologies, 15596 026) from frozen muscle samples (10 to 20 mg), quantification, and reverse transcription (iScript(tm) Reverse transcription supermix, BioRad) were performed as previously described (66). Relative mRNA expression was measured by qPCR (QuantStudio 7 Flex, Applied Biosystems, Foster City, CA) using SsoAdvanced Universal SYBR Green Supermix (Bio-Rad). Primers were designed using Primer-BLAST to include all splice variants, and were purchased from Sigma-Aldrich (Supplementary Table 4). The expression of each target gene was normalised to the geometric mean of expression of the three most stably expressed reference genes (as previously described) (66), and using the 2^−ΔΔCt^ method (where Ct is the quantification cycle). Due to insufficient muscle sample collections for 1 participant, only samples from 7 participants in the NS group were prepared for RT-PCR.

### Statistical analysis

Statistical analyses were conducted using the statistical software package GraphPad Prism (V7.03). Pre-to post-intervention changes in gene expression, protein content, mitochondrial respiratory function, peripheral skin temperature, glucose tolerance, citrate synthase activity, and plasma insulin concentrations were assessed for each group using a mixed analysis of variance (ANOVA) with one between-subjects measure (group) and a within-subjects measure (time). Significant effects of interaction (group × time), time (pre vs post), and group (NS vs SR vs SR+EX) are reported where effects are seen. Where significant effects occurred, Bonferonni post-hoc testing was performed to locate the differences. All statistical analyses of gene expression and protein content data were conducted using raw values. Gene expression data in-text are reported as percent fold-changes from pre-intervention values (mean % ± SD) with 95% confidence intervals (CI) from fold-change data (as a percentage), with the *P* value from the raw data reported. Data in figures represent fold-changes from pre-intervention values, with individual responses. A one-way ANOVA was used to assess differences between groups for the mean actigraphy sleep data and SarcPS data. All data in text, figures and tables are presented as mean ± standard deviation (SD), and 95% confidence intervals with *P* values ≤ 0.05 indicating statistical significance. Exact *P* values are presented, unless *P* < 0.001 or > 0.999.

## Additional information

### Competing interests

The authors have no competing interests to declare.

### Author contributions

NS, GR, SP, DB and JB were involved in the conception and design of the work, NS, ML, NP, AG, TS, JK, KE, ES, SP, DB and JB were involved in the acquisition, analysis or interpretation of the data for the work. NS, ML, NP, AG, GR, TS, JK, KE, ES, SP, DB and JB were involved in drafting the work and revising it critically for important intellectual content. All authors approved the final version of the manuscript. All authors agree to be accountable for all aspects of the work in ensuring that questions related to the accuracy or integrity of any part of the work are appropriately investigated and resolved. All persons designated as authors qualify for authorship, and all those who qualify for authorship are listed.

## Funding

This publication was supported in part by a Sports Medicine Australia (SMA) Research Foundation Grant and Australian Postgraduate Award PhD Scholarship to NS. SMP gratefully acknowledges support from the Canada Research Chairs program and funding for this work provided by the National Science and Engineering Research Council (NSERC) of Canada. TS was supported an NSERC of Canada graduate scholarship.

### Acknowledgements

We would like to thank all participants for their willingness to be involved with this study. We acknowledge the assistance of Dwight Marinkovic during data collection and Conor Verbruggen and Dr. Chris McGlory throughout the analysis process.

## Notes

### Competing Interest Statement

The authors have declared no competing interest.

