## Supplementary Table 1 for "Exercise mitigates sleep-loss-induced changes in glucose tolerance, mitochondrial function, sarcoplasmic protein synthesis, and circadian rhythms"

**Supplementary Data:**

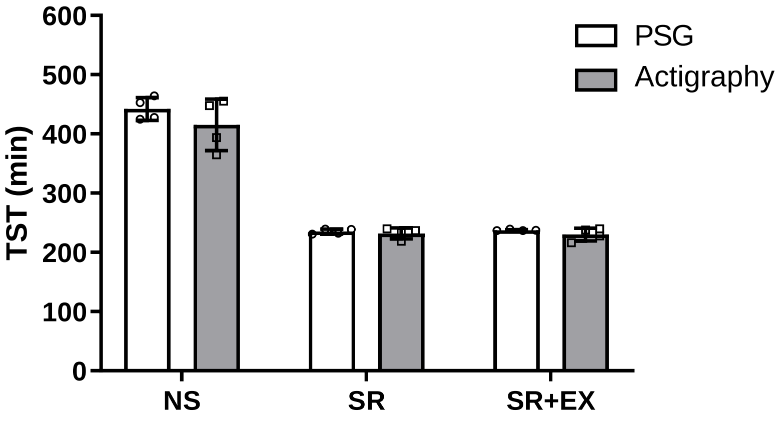


**Supplementary Figure 1 – Total sleep time (TST) measured via polysomnography (PSG) and actigraphy for night 6 of the study.** Normal Sleep (NS), Sleep Restriction (SR) and Sleep Restriction and Exercise (SR+EX), *n*=4 per group (same participants).

**Supplementary Table 1 - Polysomnography sleep analysis from night 6 of the study.**

| | NS (n=4) | SR (n=4) | SR+EX (n=4) |
|---|---|---|---|
| **TST (min)** | 442 ± 19 | 235 ± 4 # | 237 ± 1 # |
| **REM (min)** | 103 ± 14 | 65 ± 20 # | 58 ± 12 # |
| **REM (% of TST)** | 23 ± 3 | 28 ± 8 | 25 ± 5 |
| **NREM (min)** | 339 ± 20 | 170 ± 18 # | 179 ± 12 # |
| **NREM (% of TST)** | 77 ± 3 | 72 ± 8 | 75 ± 5 |
| **N1 (min)** | 27 ± 10 | 9 ± 2 # | 5 ± 2 # |
| **N1 (% of TST)** | 6 ± 2 | 4 ± 1 | 2 ± 1 # |
| **N2 (min)** | 240 ± 32 | 86 ± 34 # | 103 ± 9 # |
| **N2 (% of TST)** | 54 ± 7 | 37 ± 15 | 43 ± 4 |
| **N3 (min)** | 72 ± 17 | 75 ± 18 | 71 ± 14 |
| **N3 (% of TST)** | 16 ± 3 | 32 ± 7 # | 30 ± 6 # |
| **Wake (min)** | 18 ± 15 | 4 ± 4 | 2 ± 1 |
| **Wake (% of TST)** | 4 ± 4 | 2 ± 2 | 1 ± 1 |
| **WASO (min)** | 18 ± 15 | 4 ± 4 | 2 ± 1 |
| **Sleep efficiency (%)** | 93 ± 5 | 98 ± 2 | 99 ± 1 |
| **Sleep latency (min)** | 13 ± 15 | 1 ± 1 | 1 ± 1 |

Values are mean ± SD NS – Normal Sleep group, SR – Sleep Restriction, SR+EX – Sleep Restriction + Exercise, TST – total sleep time, REM – Rapid eye-movement, NREM – non-rapid eye-movement, N1 – nREM stage 1, N2 – nREM stage 2, N3 – nREM stage 3, WASO – Wake after sleep onset. # denotes significantly different from the NS group ($P$<0.05).

**Supplementary Table 2 - Analysis of plasma glucose (mmol.L$^{-1}$) measurements during the oral glucose tolerance test (OGTT), pre- and post-intervention**

| Time (min) | NS | | SR | | SR+EX | |
|---|---|---|---|---|---|---|
| | *Pre* | *Post* | *Pre* | *Post* | *Pre* | *Post* |
| **0** | 5.1 ± 0.4 | 5.0 ± 0.7 | 5.2 ± 0.3 | 5.0 ± 0.2 | 5.2 ± 0.2 | 5.0 ± 0.2 |
| **10** | 5.9 ± 0.8 | 6.0 ± 1.2 | 5.8 ± 0.9 | 5.7 ± 0.8 | 5.6 ± 0.3 | 5.3 ± 0.4 |
| **20** | 6.9 ± 1.2 | 7.0 ± 1.6 | 6.6 ± 0.9 | 7.2 ± 1.3 | 6.8 ± 0.6 | 6.3 ± 0.7 |
| **30** | 7.3 ± 1.9 | 6.7 ± 1.8 | 6.9 ± 1.1 | 8.5 ± 1.3* | 7.3 ± 1.0 | 7.0 ± 0.9 |
| **60** | 5.6 ± 2.1 | 4.9 ± 1.4 | 5.9 ± 1.4 | 8.4 ± 1.3* | 5.1 ± 1.2 | 6.6 ± 1.5* |
| **90** | 4.8 ± 1.8 | 4.1 ± 1.1 | 5.1 ± 1.2 | 5.9 ± 1.1 | 4.1 ± 0.9 | 5.2 ± 0.9* |
| **120** | 4.3 ± 1.4 | 4.0 ± 1.2 | 4.2 ± 0.9 | 5.1 ± 1.5 | 4.6 ± 0.6 | 4.7 ± 0.9 |
| **Mean glucose** | 5.7 ± 1.2 | 5.4 ± 1.1 | 5.7 ±0.2 | 6.5 ± 0.4* | 5.5 ± 0.3 | 5.7 ± 0.4 |
| **Total AUC** | 676.5 ± 188.6 | 617.3 ± 136.0 | 678.4 ± 32.4 | 827.2 ± 55.5* | 638.4 ± 49.8 | 705.4 ± 49.8 |

Values are mean ± SD mmol.L$^{-1}$. Area under the curve (AUC), Normal Sleep (NS), Sleep Restriction (SR) and Sleep Restriction + Exercise (SR+EX). *Denotes significant effect from pre-intervention ($P$<0.05). n=8 per group.

**Supplementary Table 3 - Analysis of plasma insulin (µIU/mL) measurements during the oral glucose tolerance test (OGTT), pre- and post-intervention**

| Time (min) | NS | | SR | | SR+EX | |
|---|---|---|---|---|---|---|
| | *Pre* | *Post* | *Pre* | *Post* | *Pre* | *Post* |
| **0** | 8.3 ± 3.1 | 8.3 ± 3.1 | 9.4 ± 3.0 | 8.7 ± 3.2 | 6.4 ± 2.8 | 6.9 ± 3.2 |
| **10** | 26.5 ± 11.7 | 31.7 ± 17.0 | 19.5 ± 8.3 | 19.0 ± 9.6 | 13.1 ± 6.3 | 14.5 ± 9.4 |
| **20** | 52.6 ± 19.7 | 75.8 ± 36.4 | 40.2 ± 16.4 | 36.4 ± 16.5 | 28.3 ± 16.2 | 26.2 ± 17.7 |
| **30** | 71.1 ± 37.4 | 83.9 ± 53.7 | 43.4 ± 19.6 | 51.3 ± 14.7 | 51.9 ± 41.3 | 43.4 ± 49.5 |
| **60** | 58.1 ± 50.5 | 43.4 ± 20.7 | 47.9 ± 32.3 | 73.5 ± 34.3* | 25.8 ± 14.0 | 35.7 ± 14.3 |
| **90** | 46.4 ± 29.1 | 27.7 ± 10.6 | 39.7 ± 28.2 | 46.2 ± 20.6 | 19.3 ± 12.9 | 30.3 ± 12.5 |
| **120** | 30.4 ±21.8 | 22.1 ± 12.4 | 21.0 ± 11.4 | 34.3 ± 21.1 | 17.1 ± 7.3 | 21.6 ± 10.2 |
| **Mean insulin** | 41.9 ± 22.4 | 42.0 ± 17.3 | 31.6 ± 2233.0 | 38.5± 1983.1 | 23.1 ± 1766.5 | 25.5 ± 1854.6 |
| **Total AUC** | 5845.3 ± 3548 | 5264.3 ± 2072 | 4454.6 ± 1958 | 5729.9 ± 1788 | 3095.7 ± 1556 | 3614.5 ± 1670 |

Values are mean ± SD µIU/mL. Area under the curve (AUC), Normal Sleep (NS), Sleep Restriction (SR) and Sleep Restriction + Exercise (SR+EX). *Denotes significant effect from pre-intervention ($P$<0.05). n=8 per group.

**Supplementary Table 4.** RT-PCR primer sequences.

| Primer Name | Primer Sequence | Product Size (bp) | Efficiency (%) | Accession No. |
|---|---|---|---|---|
| **Target Genes** | | | | |
| ***p53*** | **F –** GTTCCGAGAGCTGAATGAGG<br>**R –** TTATGGCGGGAGGTAGACTG | 123 | 101.8 | NM_001126118.1 |
| ***Tfam*** | **F –** CCGAGGTGGTTTTCATCTGT<br>**R –** GCATCTGGGTTCTGAGCTTT | 110 | 111.1 | NM_003201.3 |
| ***Dnm1l*** | **F –** CACCCGGAGACCTCTCATTC<br>**R –** CCCCATTCTTCTGCTTCCAC | 138 | 116.4 | NM_001330380.1 |
| ***Nrf2*** | **F –** AAGTGACAAGATGGGCTGCT<br>**R –** TGGACCACTGTATGGGATCA | 87 | 92.1 | NM_001197297.1 |
| ***Nrf1*** | **F –** CTACTCGTGTGGGACAGCAA<br>**R –** AGCAGACTCCAGGTCTTCCA | 143 | 102.5 | NM_005011.5 |
| ***Glut4*** | **F –** CTTCATCATTGGCATGGGTTT<br>**R –** AGGACCGCAAATAGAAGGAAGA | 75 | 85.4 | NM_001042.2 |
| ***Pgc-1α total*** | **F –** CAG CCT CTT TGC CCA GAT CTT<br>**R –** TCACTGCACCACTTGAGTCCAC | 101 | 103.6 | NM_013261.3 |
| ***β-Had*** | **F –** TGGACAAGTTTGCTGCTGAACAT<br>**R –** TTTCATGACAGGCACTGGGT | 137 | 80.6 | NM_001184705.2 |
| ***Pdk4*** | **F –** GCAGCTACTGGACTTTGGTT<br>**R –** GCGAGTCTCACAGGCAATTC | 84 | 99.7 | NM_002612.4 |
| ***Ampkα*** | **F –** CAGGGACTGCTACTCCACAGAGA<br>**R –** CCTTGAGCCTCAGCATCTGAA | 85 | 76.8 | NM_001355028.1 |
| ***Mfn1*** | **F –** CAGAAAGTGGTGTGGCACTTG<br>**R –** TTTCACTGCTGACTGCGAGAT | 104 | 112 | NM_033540.2 |
| ***Mfn2*** | **F –** CCCCCTTGTCTTTATGCTGATGTT<br>**R –** TTTTGGGAGAGGTGTTGCTTATTTC | 168 | 140 | NM_014874.3 |
| ***Pink1*** | **F –** TGTGGAACATCTCGGCAGGT<br>**R –** AGTGACTGCTCCATACTCCC | 110 | 101.8 | NM_032409.2 |
| ***Park2*** | **F –** CACGACCCTCAACTTGGCTA<br>**R –** GGTACCGGTTGTACTGCTCT | 112 | 103.4 | NM_013988.2 |
| ***Bmal1*** | **F –** GCACGACGTTCTTTCTTCTGT<br>**R –** GCAGAAGCTTTTTCGATCTGCTTTT | 114 | 97.1 | NM_001351820.1 |
| ***Clock*** | F – CGTCTCAGACCCTTCCTCAAC<br>R – GTAAATGCTGCCTGGGTGGA | 71 | 92.5 | NM_001267843.1 |
| ***Cry1*** | **F –** ACTGCTATTGCCCTGTTGGT<br>**R –** GACAGGCAAATAACGCCTGA | 74 | 104.7 | NM_004075.4 |
| ***Per1*** | **F –** ATTCGGGTTACGAAGCTCCC<br>**R –** GGCAGCCCTTTCATCCACAT | 101 | 94.8 | NM_002616 |
| ***Per2*** | **F –** CATGTGCAGTGGAGCAGATTC<br>**R –** GGGGTGGTAGCGGATTTCAT | 109 | 93.8 | NM_022817 |
| ***Rev-erb α*** | **F –** ACAGATGTCAGCAATGTCGC<br>**R –** CGACCAAACCGAACAGCATC | 73 | 102.2 | NM_005126 |
| **Housekeeping Genes** | | | | |
| ***B2M*** | **F –** TGCTGTCTCCATGTTTGATGTATCT<br>**R –** TCTCTGCTCCCCACCTCTAAGT | 86 | 98 | NM_004048.2 |
| ***ACTB*** | **F –** GAGCACAGAGCCTCGCCTTT<br>**R –** TCATCATCCATGGTGAGCTGGC | 70 | 107 | NM_001101.3 |
| ***TBP*** | **F –** CAGTGACCCAGCAGCATCACT<br>**R –** AGGCCAAGCCCTGAGCGTAA | 205 | 99 | NM_003194.4 |

**F** - Forward primer, **R** - Reverse primer
